# YAP/TAZ Signaling in Endothelial Cells Mediates the Pathogenesis of Abdominal Aortic Aneurysm Formation

**DOI:** 10.64898/2026.07.01.735919

**Authors:** Walker R. Ueland, Paolo Bellotti, Jeff A. Valisno, Aravinthan Adithan, Denny J.M. Kollareth, Jonathan R. Krebs, Michael J. Fassler, Gang Su, Shiven Sharma, Xuanxuan Yu, Guoshaui Cai, Ashish K. Sharma, Gilbert R. Upchurch

## Abstract

**Background:** Abdominal aortic aneurysms (AAA) are characterized by dilation of the aorta that can lead to aortic rupture and death. The transcriptional co-activators Yes-Associated Protein (YAP) and WW-domain-containing transcriptional co-activator with PDZ-binding motif (TAZ) are mechanosensitive effectors of the highly conserved Hippo signaling pathway. It is hypothesized that cell-specific YAP/TAZ signaling in endothelial cells (EC) plays a pivotal role in mediating AAA formation and rupture.

**Methods:** Single-cell RNA-sequencing in human AAAs was performed and differentially expressed genes (DEGs) were identified in the endothelial cell cluster. YAP/TAZ mRNA and protein expression were also assessed in human AAA and control aortic tissue. Two established murine AAA models were used with male C57BL/6 and *EC-CreER^T^*^2^*-YAP^fl/fl^/TAZ^fl/fl^* mice with/without Verteporfin (VPF, YAP/TAZ inhibitor) and XMU-MP-1 (YAP/TAZ activator) treatments. On postoperative days 14 and 28, aortic diameter, histology, cytokine, and MMP2 expressions were evaluated.

**Results:** A significant alteration in EC-specific differentially expressed YAP/TAZ-related genes was observed in which 242 genes were upregulated and 71 genes were downregulated in AAA compared to controls. Human AAA tissue showed a significant increase in YAP and TAZ protein expressions compared to controls. Elastase-treated EC-YAP/TAZ^-/-^ mice showed a significant decrease in AAA diameter compared to littermate controls. Histological quantification revealed preservation of α-smooth muscle actin, reduced elastin fiber breaks, and decreased macrophage infiltration in EC-YAP/TAZ^-/-^ mice compared to littermate controls. Importantly, pharmacological inhibition of YAP/TAZ using VPF significantly attenuated AAAs in two experimental murine models. *In vitro* data demonstrates that VPF inhibits endothelial cell YAP expression, downregulating pathways associated with pathogenic angiogenesis and vascular inflammation.

**Conclusions:** These data suggest that EC-specific YAP/TAZ signaling mediates AAA formation. Pharmacological inhibition of the Hippo pathway can significantly mitigate aortic inflammation and vascular remodeling to decrease the progression of AAAs and prevent aortic rupture.

**Highlights:**

- Expression of YAP/TAZ in endothelial cells is dysregulated in human AAAs.
- Experimental murine models demonstrate that: *a)* endothelial-cell specific deletion of YAP/TAZ protects against AAA formation, and pharmacologic alteration with *b)* verteporfin attenuates AAA formation, and *c)* XMU-MP-1 significantly exacerbates aortic inflammation, vascular remodeling, and rupture.
- Verteporfin inhibits endothelial YAP/TAZ activation by modulating ECM remodeling, pathogenic angiogenesis, pro-inflammatory cytokine and chemokine expression.

## INTRODUCTION

Abdominal aortic aneurysms (AAAs) are characterized by a significant dilation of the aorta. Over time, this ballooning can lead to aortic rupture and sudden death, representing a serious medical problem with high mortality rates. AAAs account for approximately 200,000 inpatient hospital admissions per year and comprise one of the leading causes of death in men.^1–3^ Currently, there are no preventive medical therapies to decrease AAA growth as once an aneurysm develops, the risk of aortic rupture increases.^4^ Multiple risk factors are associated with the clinical development of AAAs, including increasing age, smoking history, and hypertension.^5^ While AAAs are characterized by leukocyte infiltration into the aortic wall media and adventitia through the production of matrix degrading enzymes and pro-inflammatory cytokines, the vascular endothelium also plays a pivotal role in the modulation and regulation of this inflammatory immune response.^6–8^ Disturbed or oscillatory flow leads to endothelial cell barrier dysfunction in AAA formation, initiating pro-inflammatory signaling cascades that lead to increased oxidative stress, enhanced cell permeability, and leukocyte recruitment to promote aortic remodeling.^5,9,10^

Recent studies have highlighted the role for the mechanoresponsive proteins Yes-Associated Protein (YAP) and its paralog WW-domain-containing Transcriptional co-activator with PDZ-binding motif (TAZ) in aneurysm development.^11–14^ These central effectors of the highly conserved Hippo pathway respond to stimuli, such as turbulent flow, cell stretching, and extracellular matrix stiffening to induce cell proliferation and structural remodeling.^15,16^ In AAA formation, studies to date have largely focused on the role of YAP/TAZ in the phenotypic modulation of smooth muscle cells (SMCs) to a synthetic and proinflammatory phenotype via the cGAS/STING pathway.^11,12,14^ However, the cell-specific role of YAP/TAZ functionality may be divergent as recent studies suggest that YAP/TAZ inhibition in endothelial cells and adventitial fibroblasts can mitigate inflammation, leukocyte infiltration, and atherogenesis.^17,18^ Thus, the overarching role of cell-specific YAP/TAZ during AAA formation remains to be fully elucidated.

Our group and others have previously shown that the vascular endothelium plays a major role in SMC and macrophage activation, leukocyte modulation, and cytokine activation that results in vascular remodeling during AAA formation.^19,20^ Specifically, we have shown that inhibition of endothelial cell calcium-gated ion channels, such as Pannexin 1, mitigates aortic inflammation, aortic remodeling, leukocyte infiltration, and MMP2 activity in murine models of AAA. In addition, we found that AAA patients have an increased expression of PANX1, with an observed mortality benefit in AAA patients treated with PANX1 inhibitors. Finally, i*n vitro* studies have shown the importance of endothelial-smooth muscle cell crosstalk via PANX1/ATP signaling and P2Y2 receptors in aortic aneurysm formation.^19^

In this study, we hypothesize that mechanotransduction mediated by EC-specific YAP/TAZ activation can modulate aortic inflammation and vascular remodeling during AAA formation.^21^ As central effectors of the Hippo Pathway, YAP/TAZ transcriptional activity is regulated by a series of upstream kinases MST1/2 and LATS1/2.^22^ Therefore, we further hypothesize that pharmacologic inhibition with VPF, a specific YAP/TAZ inhibitor, can mitigate AAA formation, while activation with XMU-MP-1, an MST1/2 inhibitor and downstream YAP/TAZ activator, exacerbates AAA formation and leads to aortic rupture.

## MATERIALS AND METHODS

### Data Availability Statement

Single-cell sequencing data were analyzed from publicly available data sets in the Gene Expression Omnibus under accession codes GSE226492 & GSE152583. Original data from this study are available by request and approval of research proposal by the corresponding author.

### Human Aortic Tissue and scRNA sequencing

Single cell (sc)-RNA sequencing data of human AAAs and controls was performed with Gene Expression Omnibus (GSE226492 & GSE152583) and reanalyzed via Seurat.^23^ Cell cluster annotation was performed by the FindMarkers function in Seurat to extract cluster-specific marker genes. Cell annotations were then assigned by cross-referencing these marker gene sets and their expression in published human and mouse cell atlases using PanglaoDB Database.^24^ Cell annotations were verified via expression of putative cell markers for endothelial cell clusters as previously described in analyses of single-cell RNA aorta data sets.^25,26^ Differential gene expression analysis in endothelial cell clusters between human AAA and controls was performed via the FindMarkers function in Seurat. Statistical significance was assessed using the Wilcoxon rank-sum test and differentially expressed genes were identified with adjusted P<0.05 and |fold change| >2. A list of YAP or TAZ-related genes (YTRGs) was queried via GeneCards and cross-referenced with our list of differentially expressed genes to identify differentially expressed YTRGs. Average expression of each differentially expressed YTRG was calculated via Seurat within sample and condition. Data was scaled using the ScaleData function and used as values for the Expression Heatmap.^27^ Collection of human aortic tissue was approved by the University of Florida Institutional Review Board (#IRB 201900153). Consent was obtained by all patients prior to surgery. Male aortic tissue was resected during open abdominal aortic aneurysm repair, while organ transplant donors were used for control samples.

### Quantitative Reverse Transcription PCR

Aortic tissue from AAA patient and control (organ-transplant donors) was homogenized in Trizol and mRNA extraction was performed per manufacturer’s instructions (RNeasy Mini Kit, Qiagen, Valencia, CA). cDNA was synthesized using the iScript cDNA Synthesis Kit (Bio-Rad, Hercules, CA). GAPDH was used as a positive control in addition to YAP and TAZ primers (ThermoFisher Scientific). Primers used for quantitative RT-PCR were: YAP Fwd: GCAACTCCAACCAGCAGCAACA, YAP Rev: CGCAGCCTCTCCTTCTCCATCTG, TAZ Fwd: ACCCACCCACGATGACCCCA, TAZ Rev: GCACCCTAACCCCAGGCCAC, GAPDH Fwd: TTGATGGCAACAATCTCCAC, GAPDH Rev: CGTCCCGTAGACAAAATGGT. qRT-PCR was performed and analyzed with a CFX Connect Real-Time PCR detection system (Bio-Rad CFX96 Real-Time System). Gene expression was calculated with the relative quantification method using the following equation: 2(-ΔCT), where ΔCt = (average gene of interest) – (average reference gene). Each PCR reaction was carried out in triplicate and the relative quantification of gene expression was quantified as fold change.

### Western Blot Analysis

Protein expression of Hippo pathway mediators in human aortic tissue was evaluated by western blot analysis. Human AAA and non-aneurysmal control aortic tissues were homogenized in RIPA lysis buffer supplemented with protease inhibitors (Sigma-Aldrich) at 4 °C. The homogenates were centrifuged to remove debris, and total protein concentration in the supernatant was determined using the bicinchoninic acid (BCA) protein assay. Equal amounts of protein were mixed with SDS–PAGE loading buffer containing β-mercaptoethanol, denatured, and resolved on Mini-PROTEAN TGX gradient gels (15-well format; Bio-Rad Laboratories). Following electrophoresis, proteins were transferred to polyvinylidene difluoride (PVDF) membranes using the Trans-Blot Turbo Transfer System (Bio-Rad Laboratories). Membranes were blocked with 5% non-fat skim milk prepared in Tris-buffered saline containing 0.1% Tween-20 (TBST) to minimize nonspecific binding. Blocked membranes were incubated overnight at 4 °C with primary antibodies targeting components of the Hippo signaling pathway, including YAP (Cell Signaling Technology, mouse, 1:1000), phospho-YAP (Cell Signaling Technology, rabbit, 1:1000), TAZ (Abclonal, mouse, 1:1000), phospho-TAZ (Cell Signaling Technology, rabbit, 1:1000), MST1 and MST2 (ThermoFisher Scientific, rabbit, 1:1000), phospho-MST1/2 (ThermoFisher Scientific, rabbit, 1:1000), LATS1/2 (ThermoFisher Scientific, rabbit, 1:1000), and phospho-LATS1/2 (ThermoFisher Scientific, rabbit, 1:1000). After washing with TBST, membranes were incubated with species-appropriate horseradish peroxidase (HRP)–conjugated secondary antibodies at a dilution of 1:2000. Protein bands were detected using chemiluminescence and visualized with a Bio-Rad imaging system according to the manufacturer’s instructions.

### Animals

Adult male 8 to 12-week-old C57BL/6J wild type (WT) mice were used for this study (Jackson Laboratory, Bar Harbor, ME). Mice were housed in a temperature-controlled room at 25 °C in 12-hour light-dark cycles as per institutional animal protocols. Mice were provided with drinking water and a standard chow diet ad libitum. All animal experiments followed protocol approved by the University of Florida’s Institutional Animal Care and Use Committee (IACUC no. 2024000000138).

### Murine Topical Elastase Model of AAA

Using an established murine topical elastase model, male WT C57BL/6 mice (8-12 weeks) were used, as previously described.^19,28^ Following aortic aneurysm induction, separate groups of C57BL/6 mice were intraperitoneally injected with either verteporfin (VPF, a specific YAP/TAZ inhibitor, 50mg/kg, Med Chem Express, Monmouth Junction, NJ), XMU-MP-1 (an MST1 kinase inhibitor, 3mg/kg, Med Chem Express, Monmouth Junction, NJ), or vehicle control (0.2 ml saline) from the day after surgery through postoperative day 13 via every other day injection. Dosage selection was based on current published literature.^18,29^ Separate groups of animals were used with *EC*-*Cre*ER*^T2^*-YAP/TAZ^fl/fl^ mice (Taconic Biosciences, Germantown, NY) that were injected with 20 mg/ml intraperitoneal tamoxifen for five days. Successful knockout was determined by qRT-PCR via murine tail clippings and confirmed by flow cytometry. Two weeks after i.p. injections, mice were anesthetized with isoflurane and underwent exposure of the infrarenal abdominal aorta as previously described.^19,28^ Briefly, the aorta was circumferentially dissected from surrounding tissues and exposed to 5 µL of peri-adventitial elastase (0.4 U/mL type 1 porcine pancreatic elastase, Sigma Aldrich, St. Louis, MO) or heat-inactivated elastase as control.

Murine aortas were harvested on day 14 and mice were euthanized under anesthesia by exsanguination. The abdominal aorta from below the left renal vein to the aortic bifurcation was dissected and exposed. The maximal aortic adventitial diameter was measured and compared to the intact self-control portion above the left renal artery using video microscopy with NIS-Elements D.5.10.01 software (Nikon SMZ-25, Melville, NY).^30^ Change in aortic diameter dilation was calculated using the formula: (maximal AAA diameter – self-control aortic diameter) / (maximal AAA diameter) x 100%. Aortic sections were harvested and snap-frozen in liquid nitrogen before storing at -80 °C or preserved in formalin or Optimal Cutting Temperature (OCT) compound for immunohistochemistry.

### Chronic AAA and Aortic Rupture Model

A second chronic AAA and aortic rupture murine model that exhibits thrombus formation was also used.^31^ Mice were given drinking water containing 0.2% β-aminopropionitrile (BAPN) two days prior to topical elastase induction until aortic harvest. Male C57BL/6 mice were induced with 5 µL of peri-adventitial elastase (0.4 U/mL type 1 porcine pancreatic elastase, Sigma Aldrich, St. Louis, MO) or heat-inactivated elastase as control. Separate groups of mice were treated with either verteporfin (VPF, 50mg/kg, Med Chem Express, Monmouth Junction, NJ), XMU-MP-1 (3mg/kg, Med Chem Express, Monmouth Junction, NJ), or vehicle control (0.2 ml saline) from postoperative day 14 through postoperative day 27. Animals were euthanized on postoperative day 28, and aortic tissue was harvested as described above.

### Histology

Aortic tissue was fixed in 4% buffered formaldehyde for 24 hours and embedded in paraffin or OCT and sectioned at 5 µm. Immunostaining was performed for α-SMA (α-smooth muscle actin), elastin (Verhoeff-van Gieson), and macrophages as previously described.^31^ Antibodies for immunohistochemical staining were anti-mouse α-SMA (1:5000, catalog no. A5691; Sigma, St. Louis, MO) and anti-mouse Mac2 (galectin-3) for macrophages (1:10000, catalog no. CL8942AP; Cedarlane Laboratories, Burlington, Ontario, Canada). Heat-inactivated elastase-treated aortic tissue was used as a positive control, while isotype IgG2a control antibody was used as a negative control for immunostaining. Histological analysis was performed on different sections of aortic tissue from each animal and quantified. Images were acquired with 20x magnification via an Olympus microscope equipped with a digital camera using NIS-Elements D.5.10.01 software (Nikon SMZ-25, Melville, NY).

Representative images from each treatment group are shown for accuracy. For histological grading, the positive staining area of the aortic media (α-SMA, VVG) and entire aorta (macrophage) was selected and measured using the integrated optical density of each section.^32^ The histological expression was then quantified with QuPath (v0.6.0) and ImageJ bioimage analysis software by importing brightfield-stained images and annotating the tissue manually.^33^ Positive cell detection was assigned by optical density to identify positive regions of interest. The percentage of positive staining area relative to the total annotated area was then calculated and reported as positively stained % area. Elastin breaks were calculated per mm^2^ aortic media tissue using ImageJ software as previously described.^34^ Quantification and analysis were applied consistently across all aortic samples.

### Cytokine Multiplex Assay

Cytokine levels in murine aortic tissue homogenates were quantified using the Bio-Plex Bead Array system with a multiplex cytokine panel assay (Bio-Rad Laboratories, Hercules, CA), following the manufacturer’s instructions. Briefly, aortic tissues from various experimental groups were snap-frozen in liquid nitrogen, ground using a mortar and pestle, and homogenized in tissue extraction buffer. Protein concentration was determined using the BCA assay kit (Thermo Fisher Scientific), and equal amounts of protein were used for cytokine measurements in the tissue samples.

### Gelatin Zymography for MMP Activity

MMP2 and MMP9 expression was measured in murine aortic tissue as previously described.^35^ Briefly, samples were loaded onto a zymogram gel (ThermoFisher Scientific) with 3 µg of isolated protein per lane. Protein samples were separated by gel electrophoresis and then renatured with buffer (ThermoFisher Scientific) for 30 minutes. The resulting gel was placed in a developing buffer and stained with SimplyBlue SafeStain (ThermoFisher Scientific). Bands were assessed by Bio-Rad Image Lab 4.0 software.

### Cell Culture and *In Vitro* Experiments

Primary human aortic endothelial cells (HAECs; Lifeline Cell Technology, Frederick, MD) were used for *in vitro* experiments. Cells were first exposed to porcine elastase (Worthington Biochemical Corporation, Lakewood, NJ) at a concentration of 0.4 U/mL for 5 minutes, as previously described.^36^ Following elastase exposure, cells were washed twice with sterile 1× phosphate-buffered saline (PBS), treated with or without verteporfin (0.35μM) and XMU-MP-1 (5μM), and then incubated in basal culture medium for 12 hours containing a cytokine mixture (cytomix) consisting of interleukin-17 (IL-17A), high mobility group box 1 (HMGB1), and tumor necrosis factor-α (TNF-α), each at a final concentration of 50 ng/mL. At the end of the treatment period, conditioned media were collected and stored at -80 °C for downstream cytokine analysis. Total cellular proteins were extracted using RIPA lysis buffer supplemented with protease and phosphatase inhibitors. Protein concentrations were quantified using the bicinchoninic acid (BCA) protein assay (ThermoFisher Scientific, Waltham, MA) according to the manufacturer’s instructions.

### Flow Cytometry

Single-cell suspensions were prepared from freshly isolated aortic tissues for flow cytometric analysis. Tissues were minced into small fragments and enzymatically digested in plain RPMI medium containing collagenase type I (2.5 mg/mL) and elastase (0.5 mg/mL) for 40 minutes at 37°C with vigorous shaking. Intermittent mechanical dissociation was performed by gentle pipetting to facilitate tissue disaggregation. Enzymatic digestion was quenched by the addition of fetal bovine serum (FBS) to a final concentration of 20% (v/v). The resulting suspension was passed through a 70-µm cell strainer to remove undigested debris, and single cells were pelleted by centrifugation at 400 × g for 5 minutes at 4°C. Cell pellets were washed twice with ice-cold 1× phosphate-buffered saline (PBS). For viability discrimination, resuspended cells were incubated with Live/Dead Fixable Blue viability dye in 1× PBS for 30 minutes at room temperature in the dark. Surface immunostaining was subsequently performed in flow cytometry staining buffer using fluorophore-conjugated antibodies against CD45 (BV785) to identify hematopoietic cells, and VE-cadherin (CD144; RB780) as an endothelial lineage marker.

For intracellular detection of YAP/TAZ, cells were fixed using BD Cytofix/Cytoperm IC Fixation Buffer according to the manufacturer’s instructions. Permeabilization and all subsequent wash steps were performed using BD Permeabilization and Wash Buffer (1×). Intracellular staining was carried out using an Alexa Fluor 647 (AF647)-conjugated anti-YAP/TAZ antibody diluted in BD Permeabilization and Wash Buffer. An isotype-matched control antibody was included in all panels to establish nonspecific background fluorescence thresholds. Following antibody incubation, cells were washed with BD Permeabilization and Wash Buffer and subsequently resuspended in flow cytometry staining buffer for data acquisition on a Cytek Aurora 5-laser spectral flow cytometer. Data was analyzed using FlowJo software (v10.10; BD Biosciences).

### *In vitro* bulk RNA sequencing

Primary human aortic endothelial cells (ATCC, Manassas, VA) were used for bulk RNA sequencing after respective treatments. Quality control (QC) on the reads was conducted by using fastqc (v0.12.1) and all reads are of 151 bps. The first 15 bases from 5’ end and the bases with score < 20 were trimmed using cutadapt (v3.4), resulting a uniformed length of 120 bps. Reads were then mapped to the human reference transcriptome (GrCh38; 10x Genomics reference version 2024-A) using star aligner (v2.7.3a).^37^ Gene expression was quantified by using RSEM, generating read count matrices with 14,051 genes for downstream analysis.^38^

Further data processing was performed on the read count matrices primarily in R v4.5.3. Differential gene expression analysis was performed using edgeR package (v4.6.3).^39^ Difference in library sizes from the raw gene count matrices were normalized using the weighted trimmed mean of M-values (TMM) method.^40^ Groups were treated as categorical variables, with the control group being the reference condition. Normalized count data was generated by adjusting for the dispersions using negative binomial likelihood. Generalized linear models (GLMs) were then fitted with the comparisons between groups assessed using likelihood ratio tests with predefined contrasts. The significant differentially expressed genes (DEGs) were defined as those with FDR p values < 0.05 and absolute log2FC greater than 1.

Enriched canonical pathways were identified using Gene Set Enrichment Analysis (GSEA). GSEA was conducted using clusterProfiler package (v4.18.4) with Hallmark and KEGG legacy gene sets from Human Molecular Signatures Database (MSigDB) as the reference database.^41,42^ For each comparison, genes were first ranked by the log fold change (log2FC) from the differential expression analysis. Pathways with statistically significant enrichment were defined with false discovery rate (FDR) < 0.05. The sign of Normalized Enrichment Score (NES) indicates the direction of regulation, where positive values indicate the genes in the particular pathway were upregulated in compared group and negative values indicate downregulation.

### Statistical Analysis

Data from all experimental results are expressed as mean±SEM. Data distribution was determined by using a Shapiro-Wilk normality test. Multiple group comparisons were made using 1-way ANOVA with Tukey post hoc analysis if they passed a normality test, or Kruskal-Wallis 1-way ANOVA if the normality test failed. Student t test or Mann-Whitney U test rank-sum test were used to compare means between two groups, depending on the normality test results. All data were analyzed using GraphPad Prism program (Version 10), and p≤0.05 was considered statistically significant.

## RESULTS

### Single-cell RNA-sequencing analysis reveals dysregulation of endothelial cell YAP/TAZ-related genes in AAA patients

To evaluate the EC-specific dysregulation in human AAA tissue, we analyzed human scRNA datasets using Seurat (Figure 1A-B). Differential expression analysis comparing ECs from human AAAs (n=7) vs. control aortas (n=5) was performed via FindMarkers calculated using Wilcoxon Rank Sum Testing. Genes with an adjusted p-value <0.05 and |fold-change| > 2 were considered significant differentially expressed genes (DEGs). A list of YAP and TAZ-related genes (YTRGs) was then generated using GeneCards and used to identify differentially expressed YTRGs (DE YTRGs). A total of 11,848 genes were detected within the EC cluster and tested for differential expression (Figure 1C). Of these differentially expressed genes, 2,534 genes were upregulated, 667 were downregulated, and 8,647 genes were unchanged (Supplementary Table S1). Furthermore, 242 of 2,534 upregulated and 71 of 667 downregulated genes were YAP or TAZ-related (Figure 1D-F). Key upregulated genes included Fos-related antigen 1 (FOS1), Serpin Family E Member 1 (SERPINE1), Lymphoid enhancer-binding factor 1 (LEF1), and C-X-C chemokine receptor type 4 (CXCR4), genes involved in endothelial activation, vascular fibrosis, extracellular matrix remodeling, and inflammation (Supplementary Table S1). To evaluate YAP/TAZ expression in human AAAs, we analyzed aortic tissue from healthy control and AAA patients. mRNA expression of TAZ was significantly increased in AAA compared to healthy controls (Supplementary Figure S1A). Total YAP and TAZ protein expression were also significantly increased in AAA compared to healthy controls (Total YAP: 4.1±1.0 vs. 0.9±0.2; n=6-7/group; p<0.05; Total TAZ: 3.9±0.9 vs. 0.4±0.1; n=6-7/group; p<0.01) (Supplementary Figure S1B-C). Protein expressions for MST1, MST2, and LATS1/2 were significantly increased in human AAA tissue compared to controls, suggesting that the Hippo pathway is deactivated in AAA formation, allowing YAP and TAZ to co-localize to the nucleus to exert downstream effects (Supplementary Figure S2A-C). Given these findings that highlighted EC-specific YAP/TAZ expression to be dysregulated in AAAs, we then explored the role of this pathway in murine models of AAA.

**Figure 1.**
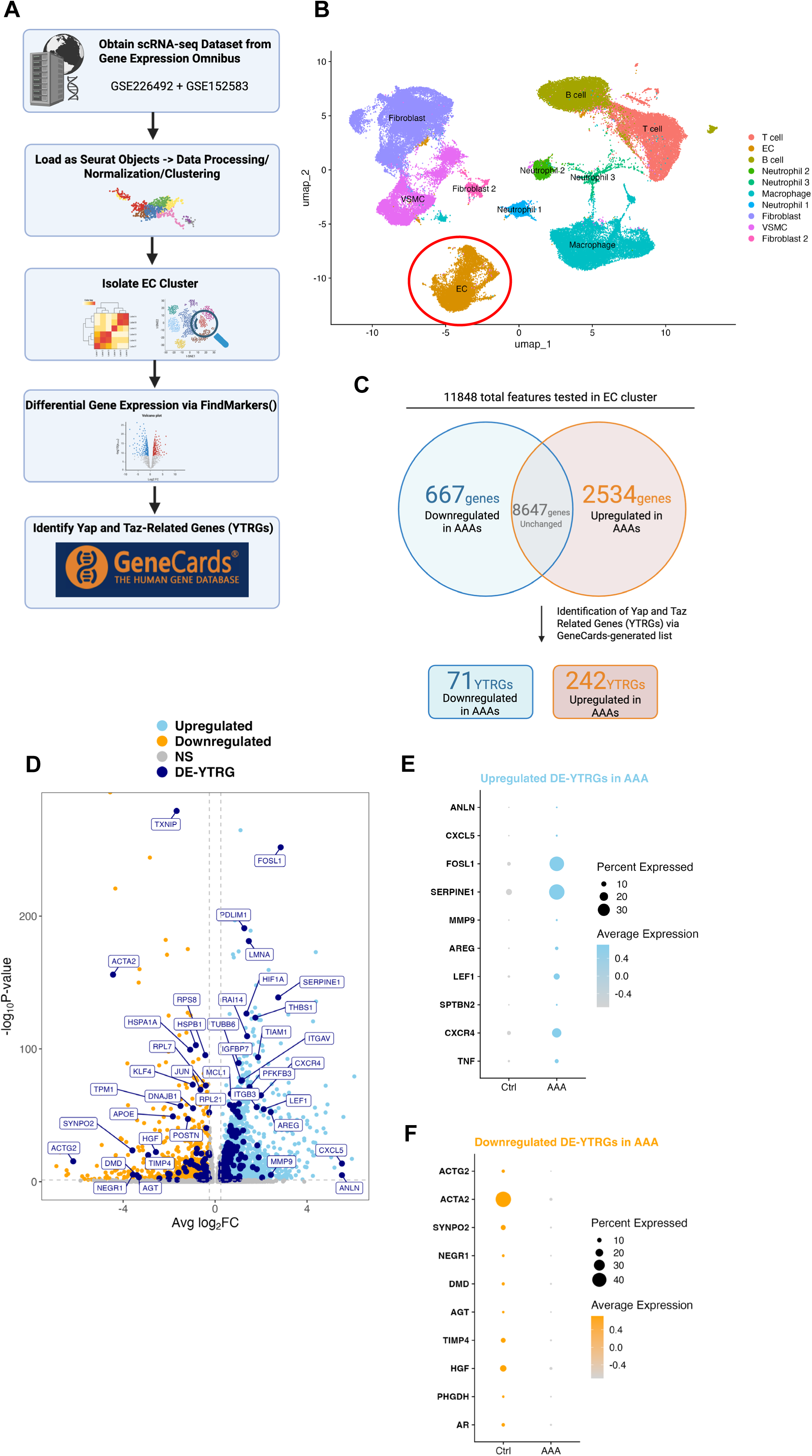
YAP and TAZ related genes are dysregulated in ECs of human AAAs. **A,** Bioinformatic workflow to re-analyze publicly available scRNA-seq data sets (GSE226492 & GSE152583) from Gene Expression Omnibus. Data was analyzed via Seurat and the EC cluster was isolated. Differential expression analysis comparing ECs from human AAAs (n=7) vs. control aortas (n=5) was performed via FindMarkers calculated using Wilcoxon Rank Sum Testing. Genes with an adjusted p-value < 0.05 and |fold-change| > 0.1 were considered significant differentially expressed genes (DEGs). A list of YAP and TAZ-related genes (YTRGs) were generated using GeneCards and used to identify differentially expressed YTRGs (DE YTRGs). **B,** Uniform manifold projection (UMAP) plot of the annotated clusters from human AAAs and control. The EC cluster is circled and subsetted for further analysis. **C,** Volcano plot displaying significantly downregulated (sky blue) and upregulated (orange) genes. DE YTRGs are highlighted and colored (navy blue). **D,** Venn diagram displaying gene counts from this analysis. A total of 11,848 genes were detected within the EC cluster and tested for differential expression. Of the 11,848 genes, 667 genes were downregulated, 2,534 were upregulated, and 8,647 were unchanged. 71/667 downregulated genes and 242/2,534 upregulated genes were YTRGs. **E-F,** DotPlot of upregulated and downregulated DE-YTRGs in ECs displayed as average expression aggregated between conditions (control vs. AAA).

### Deletion of YAP/TAZ in ECs mitigates AAA formation

Using the murine elastase AAA model, we observed that aortic diameter sequentially increased in WT mice over time (Day 14: 169±10.5% and Day 7: 105±5.6% compared to Day 0: 6±0.5%; n=5-10/group; p<0.0001). Also, concurrently YAP/TAZ expression in ECs significantly increased over time (Day 14: 64±1.9% and Day 7: 75±0.5% vs. Day 0: 41±2%; n=3/group; p<0.001) (Supplemental Figure S3A-C).

As EC-specific YAP/TAZ expression increased over time throughout murine AAA formation, we next generated *EC-CreER^T2^-YAP^fl/fl^/TAZ^fl/fl^*transgenic mice and used the topical elastase AAA model (Figure 2A). EC-specific YAP/TAZ deletion was confirmed by flow cytometry using murine aortic samples (Supplementary Figure S4). EC-YAP/TAZ^-/-^ mice exhibited a significant decrease in AAA diameter compared to EC-YAP/TAZ^+/+^ littermate controls (111.8±7.8 vs. 161.4±11.5; n=13-14; p<0.001; Figure 2B-C). Of note, there was no significant difference in aortic diameter between elastase-treated wild type (WT) mice and EC-YAP/TAZ^+/+^ littermate controls (156.6±5.0 vs. 161.4±11.5; n=13-15).

**Figure 2.**
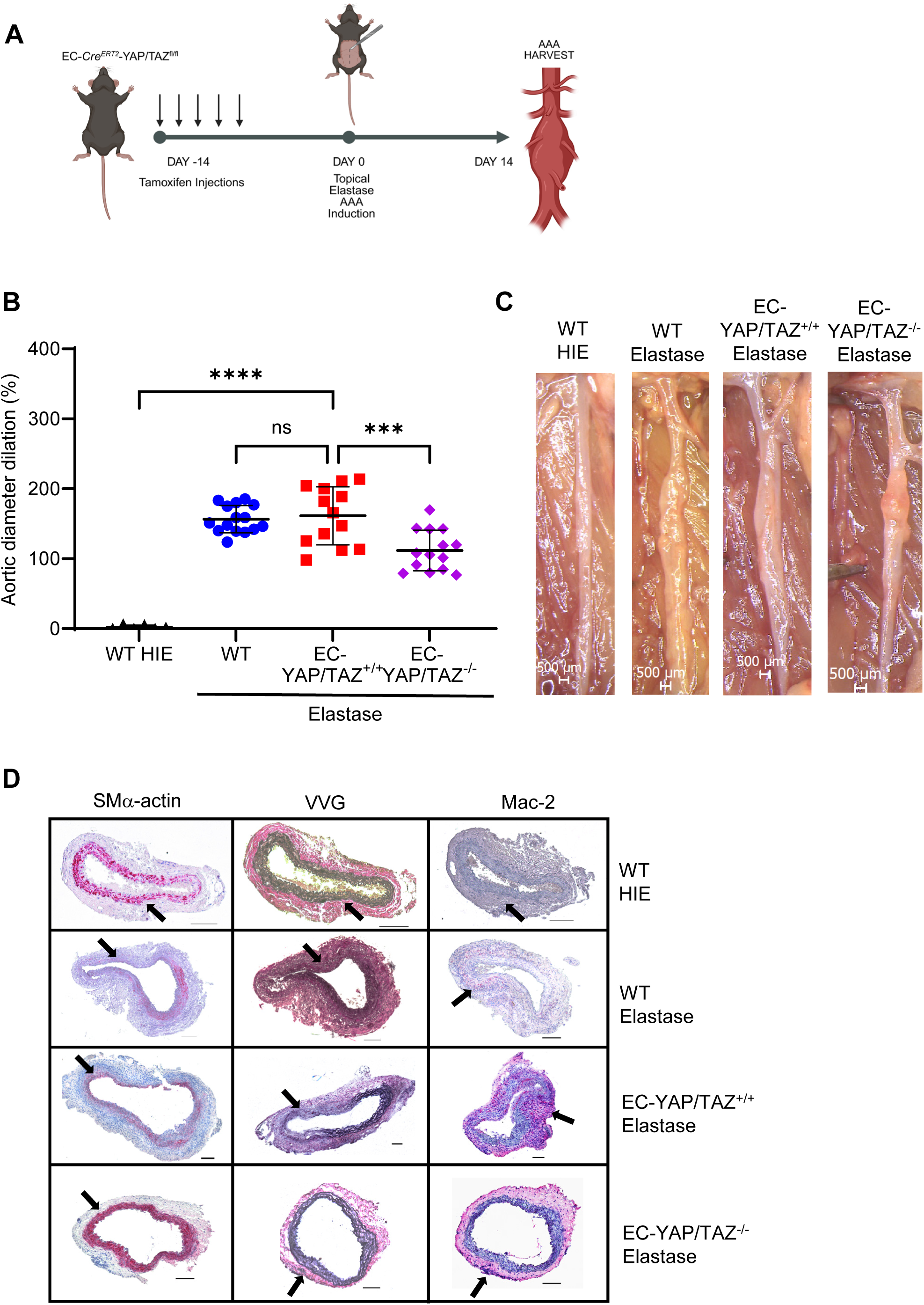
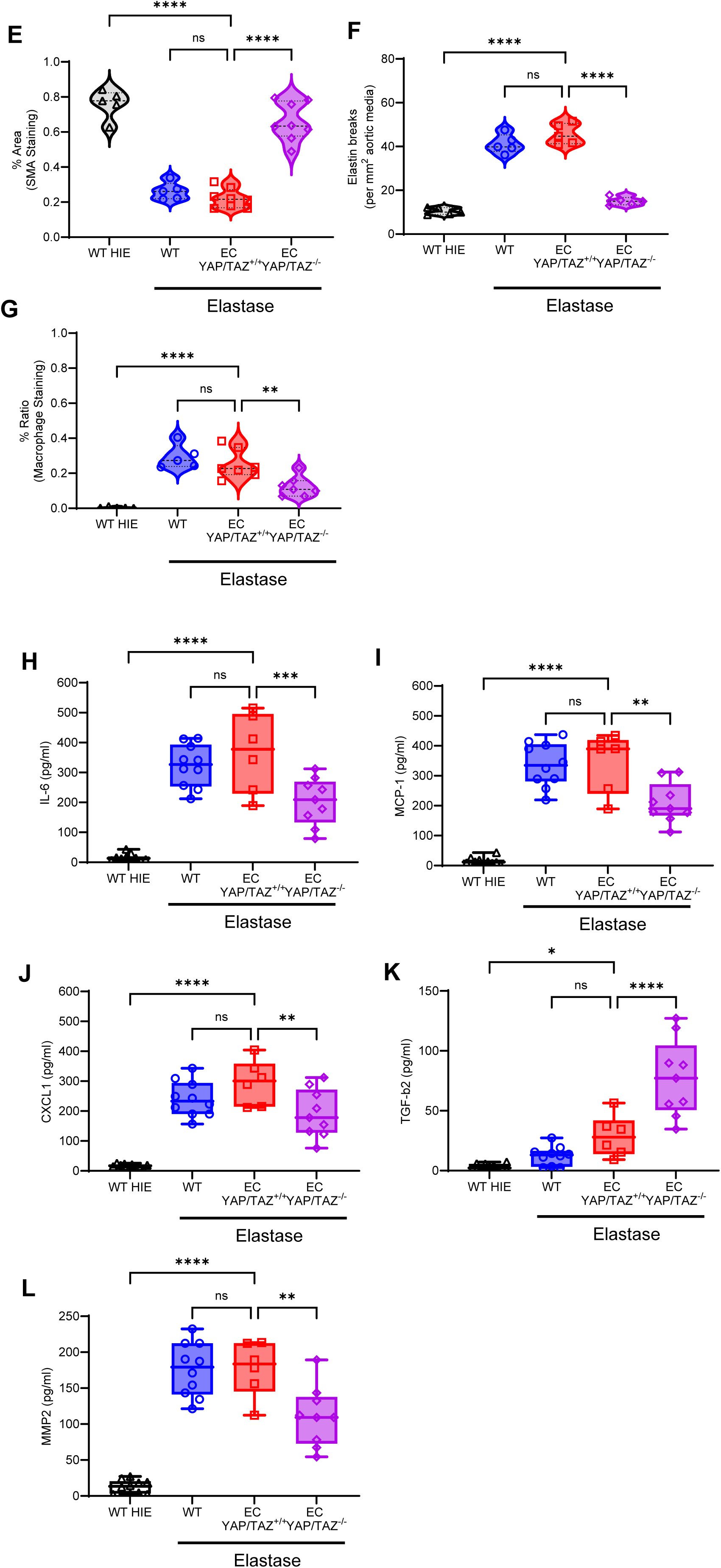
Deletion of YAP/TAZ on ECs mitigates AAA formation. **A,** Schematic representing the topical elastase AAA model. Mice were injected with tamoxifen in *EC-CreER^T2^-YAP/TAZ^fl/fl^*(EC-YAP/TAZ^-/-^) for five days to induce deletion of YAP/TAZ in ECs, followed by AAA induction with topical elastase or deactivated elastase (heat-inactivated; controls) on day 14. On postoperative day 14, abdominal aortas were measured with video micrometry and harvested for histology and cytokine analysis. **B,** EC-YAP/TAZ^-/-^ mice showed a significant reduction in aortic diameter compared to EC-YAP/TAZ^+/+^ littermate controls. ****p<0.0001, ***p<0.001; ns, not significant; n=6-14/group; 1-way ANOVA. **C,** Representative images of aortic phenotype in all groups. Scale bar= 500μM. **D-G,** EC-YAP/TAZ^-/-^ mice exhibited preserved aortic morphology as observed by preserved SM α-actin expression, decreased elastin fiber breakage, and reduced macrophage infiltration, compared to EC-YAP/TAZ^+/+^ littermate controls. Scale bar=100μM; ****p<0.0001, **p<0.01; n=5-8/group; 1-way ANOVA. **H-L,** Aortic tissue from EC-YAP/TAZ^-/-^mice treated with elastase showed a significant decrease in pro-inflammatory cytokine expression and MMP2 content compared to littermate controls. ****p<0.0001, ***p<0.001, **p<0.01, *p<0.05; n=6-9/group; 1-way ANOVA.

Comparative histology and immunostaining of aortic specimens showed preserved aortic morphology, with increased SMα-actin expression (65.9±3.9% vs. 22.2±2.0%; p<0.0001), decreased elastin fiber disruption (15.0±0.8 vs. 45.7±2.1; p<0.0001), and reduced macrophage infiltration (12.4±2.2% vs. 25.1±3.1%; p<0.01) compared to EC-YAP/TAZ^+/+^ littermate controls (Figure 2D-G). Furthermore, EC-YAP/TAZ^-/-^ mice exhibited a significant reduction in pro-inflammatory cytokines and chemokines, including IL-6, MCP-1, CXCL1, and MMP2 compared to EC-YAP/TAZ^+/+^ littermate controls (Figure 2H-L). Additionally, elastase-treated EC-YAP/TAZ^-/-^ mice showed reduced MMP2 and -9 activities compared to elastase-treated EC-YAP/TAZ^+/+^ mice on day 14 (Supplemental Figure S5). Collectively, these results demonstrate that deletion of YAP/TAZ on ECs protects against aortic inflammation, vascular remodeling and AAA formation.

### Pharmacologic inhibition of YAP/TAZ mitigates AAA formation

To investigate the effects of pharmacologically altering the YAP/TAZ signaling pathway, we used VPF (a specific YAP/TAZ inhibitor) in the C57BL/6 (WT) mice using the topical elastase AAA model (Figure 3A-B). Administration of VPF significantly attenuated aortic diameter compared to elastase-treated WT mice (81.8±6.9 vs. 142.0±4.6; n=20-24; p<0.0001) (Figure 3C-D). Conversely, activation of YAP/TAZ via XMU-MP-1 significantly increased aortic diameter size compared to elastase-treated WT mice (185.4±8.2 vs. 142.0±4.6; n=24-31; p<0.0001) (Figure 3C-D).

**Figure 3.**
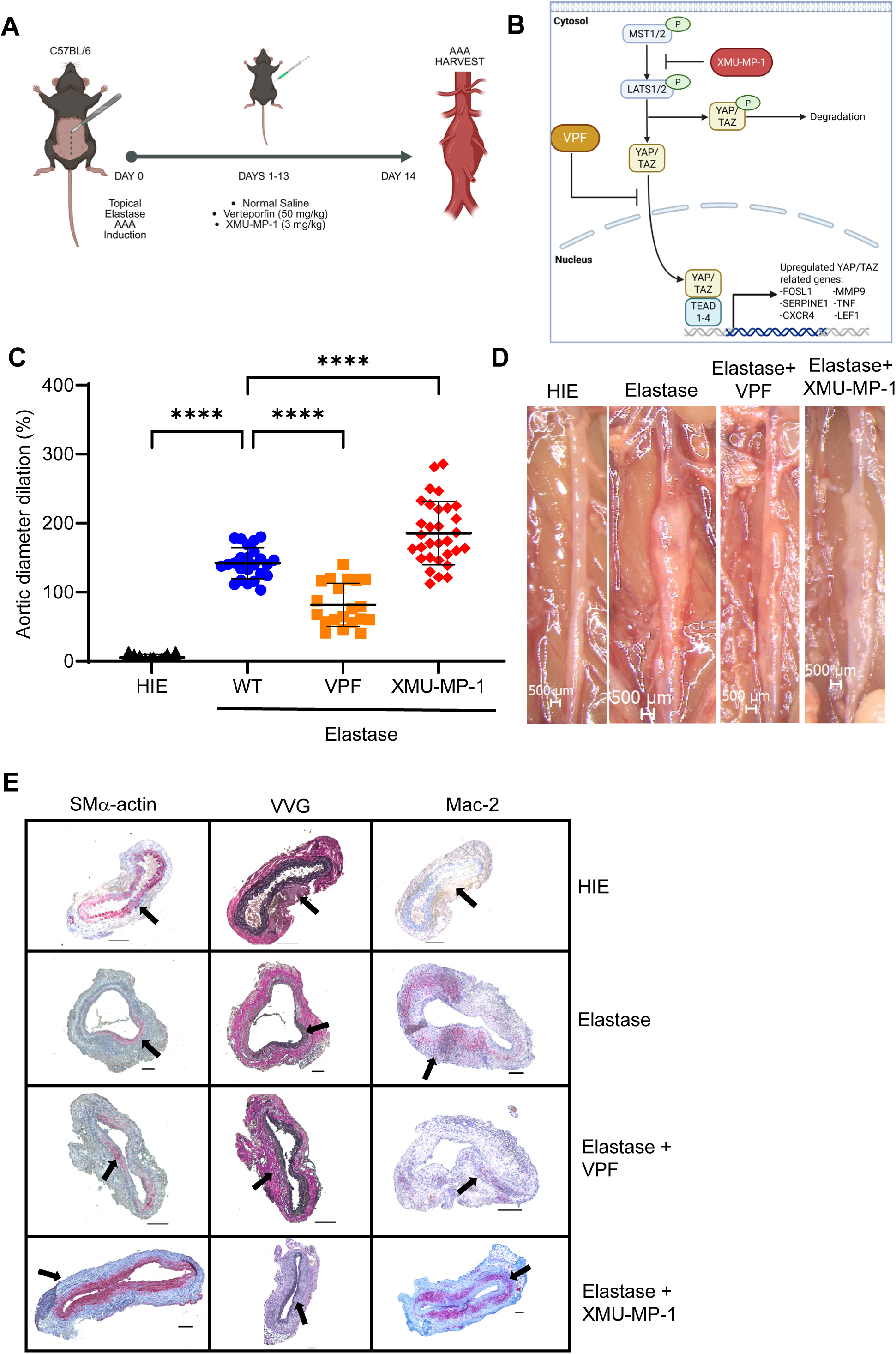
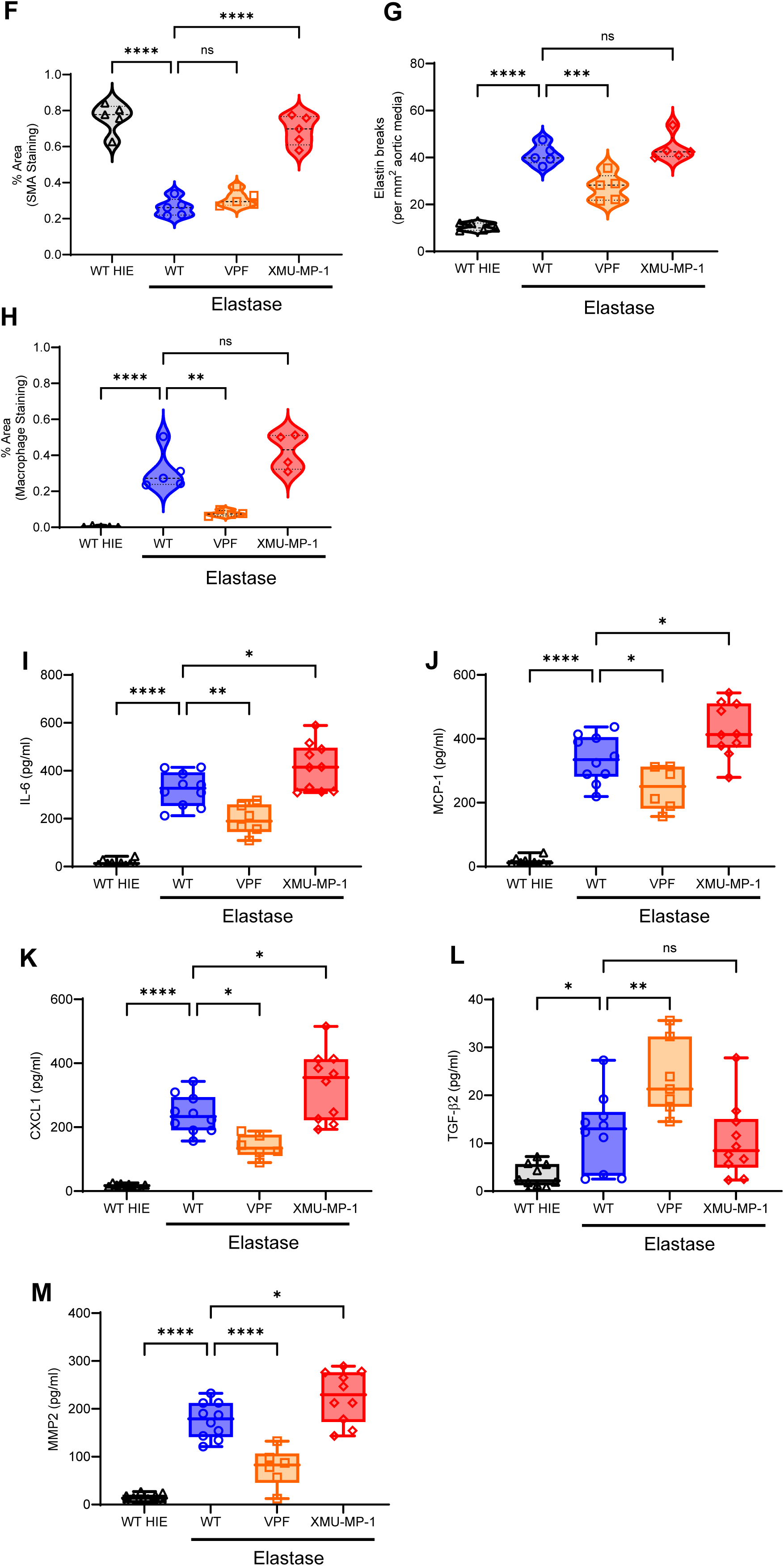
Verteporfin treatment attenuates AAA formation. **A,** Schematic description of topical elastase AAA model with/without administration of VPF (YAP/TAZ inhibitor), or XMU-MP-1 (YAP/TAZ activator). On postoperative day 14, abdominal aortas were measured with video micrometry and harvested for tissue analysis. **B**, Mechanistic signaling of pharmacological modulation targeting the Hippo pathway mediators. **C,** Verteporfin treatment significantly mitigate AAA diameter compared to elastase-treated WT mice. Conversely, XMU-MP-1 significantly increased aortic diameter compared to mice treated with topical elastase. ****p<0.0001; n=20-31/group; 1-way ANOVA. **D,** Representative images of aortic phenotype in all groups. Scale bar=500μM. **E-H,** In VPF-treated WT mice, comparative histology demonstrated no differences in SM α-actin expression, but significantly reduced elastin fiber disruption, and macrophage infiltration compared to elastase-treated mice. Conversely, XMU-MP-1 treatment preserved SM α-actin expression, but significantly increased elastin fiber disruption and macrophage infiltration. Scale bar=100μM; ****p<0.0001, ***p<0.001, **p<0.01; ns, not significant; n=5/group; 1-way ANOVA. **I-M,** Aortic tissue from WT mice treated with VPF showed a significant decrease in pro-inflammatory cytokine expression and MMP2 content compared to elastase-treated WT mice. Conversely, administration of XMU-MP-1 significantly increased pro-inflammatory cytokine and chemokine expression. ****p<0.0001, **p<0.01, *p<0.05; ns, not significant; n=6-10/group; 1-way ANOVA.

In VPF-treated mice, comparative histology and immunohistochemistry demonstrated no difference in SMα-actin staining (31.0±1.9% vs. 26.3±2.2%), but reduced elastin fiber disruption (27.3±2.6 vs. 41.2±2.0; p<0.001) and macrophage infiltration (7.7±1.0% vs. 31.3±4.9%; p<0.01) compared to elastase-treated WT mice (Figure 3E-H). Interestingly, treatment with XMU-MP-1 preserved α-SMA expression (69.0±3.7% vs. 26.3±2.2%; p<0.0001) but there was no significant difference in elastin breaks (44.0±2.5 vs. 41.2±1.9) or macrophage infiltration in XMU-MP-1-treated mice (42.2±5.1% vs. 31.3±4.9) compared to elastase-treated WT mice (Figure 3E-H). On cytokine analysis, murine aortas treated with VPF demonstrated a reduction of pro-inflammatory cytokines, including IL-6, MCP-1, CXCL1, as well as MMP2 expression, while mice treated with XMU-MP-1 demonstrated an increase in pro-inflammatory cytokine profiles compared to elastase-treated WT mice (Figure 3I-M). Together, these findings demonstrate that inhibition of YAP/TAZ with VPF can mitigate aneurysm growth, influence aortic remodeling, and decrease pro-inflammatory cytokines and chemokines, while activation via XMU-MP-1 leads to larger aneurysms and promotes inflammation.

### VPF treatment attenuates pre-formed AAA formation to protect against aortic rupture

A second chronic inflammatory elastase+BAPN model of AAA and rupture was used to further elucidate the role of pharmacologically altering the YAP/TAZ signaling pathway. C57BL/6 mice exposed to elastase+BAPN were treated with either VPF or XMU-MP-1 from postoperative days 14-27 after undergoing aortic aneurysm induction (Figure 4A). Verteporfin-treated mice exhibited a significant reduction in aortic diameter compared to elastase+BAPN-treated mice alone (185.8±13.3 vs. 248.7±12.4; n=23/group; p<0.01) (Figure 4B-C). Conversely, mice treated with XMU-MP-1 had significantly larger aortic diameters compared to elastase+BAPN-treated mice (412.8±32.3 vs. 248.7±12.4; n=12-23/group; p<.0001) (Figure 4B-C). Furthermore, XMU-MP-1-treated mice were more prone to aortic rupture and demonstrated a significant reduction in survival compared to elastase+BAPN-treated mice, with only 40% surviving to the twenty-eight-day endpoint (log-rank p=0.0132) (Figure 4D).

**Figure 4.**
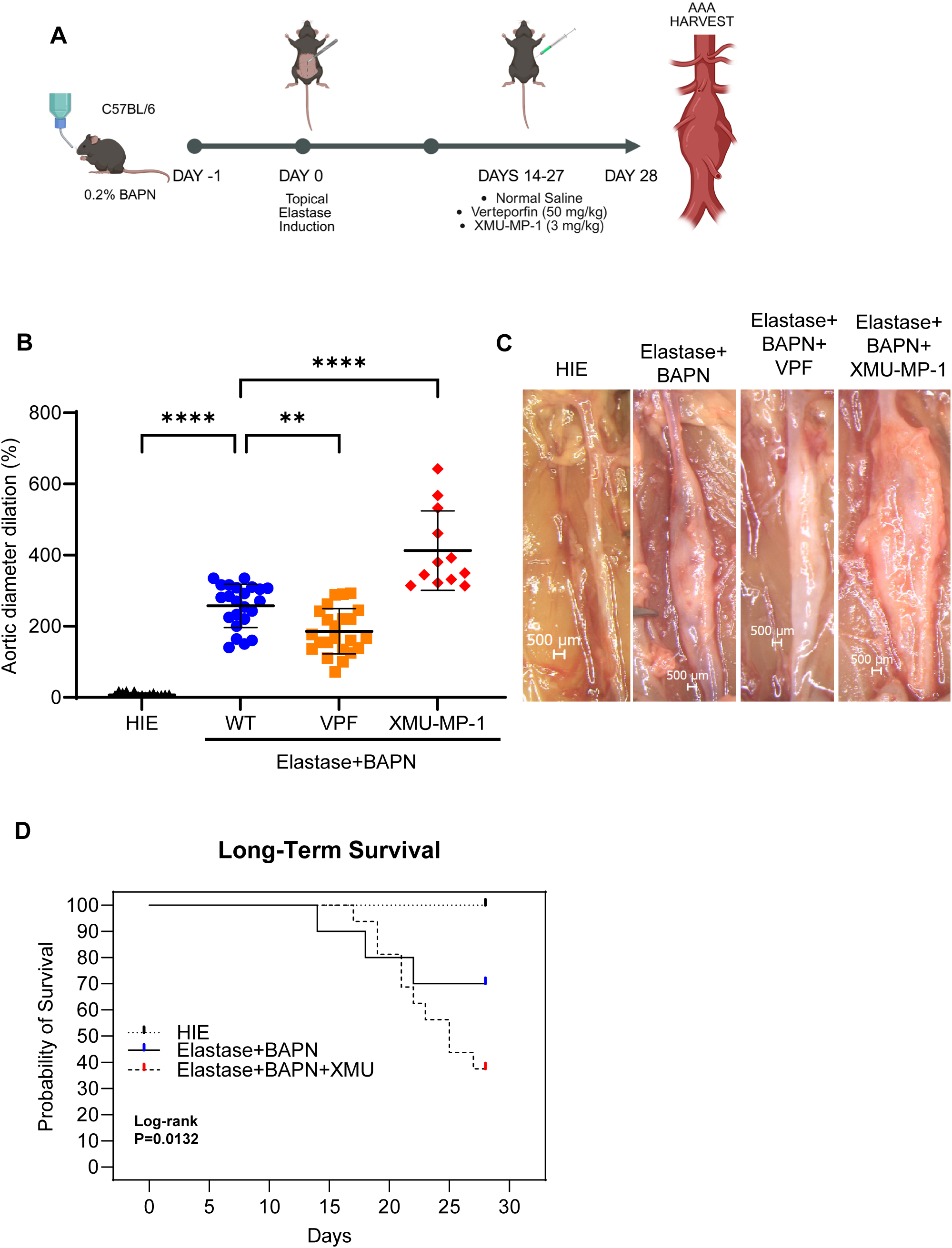
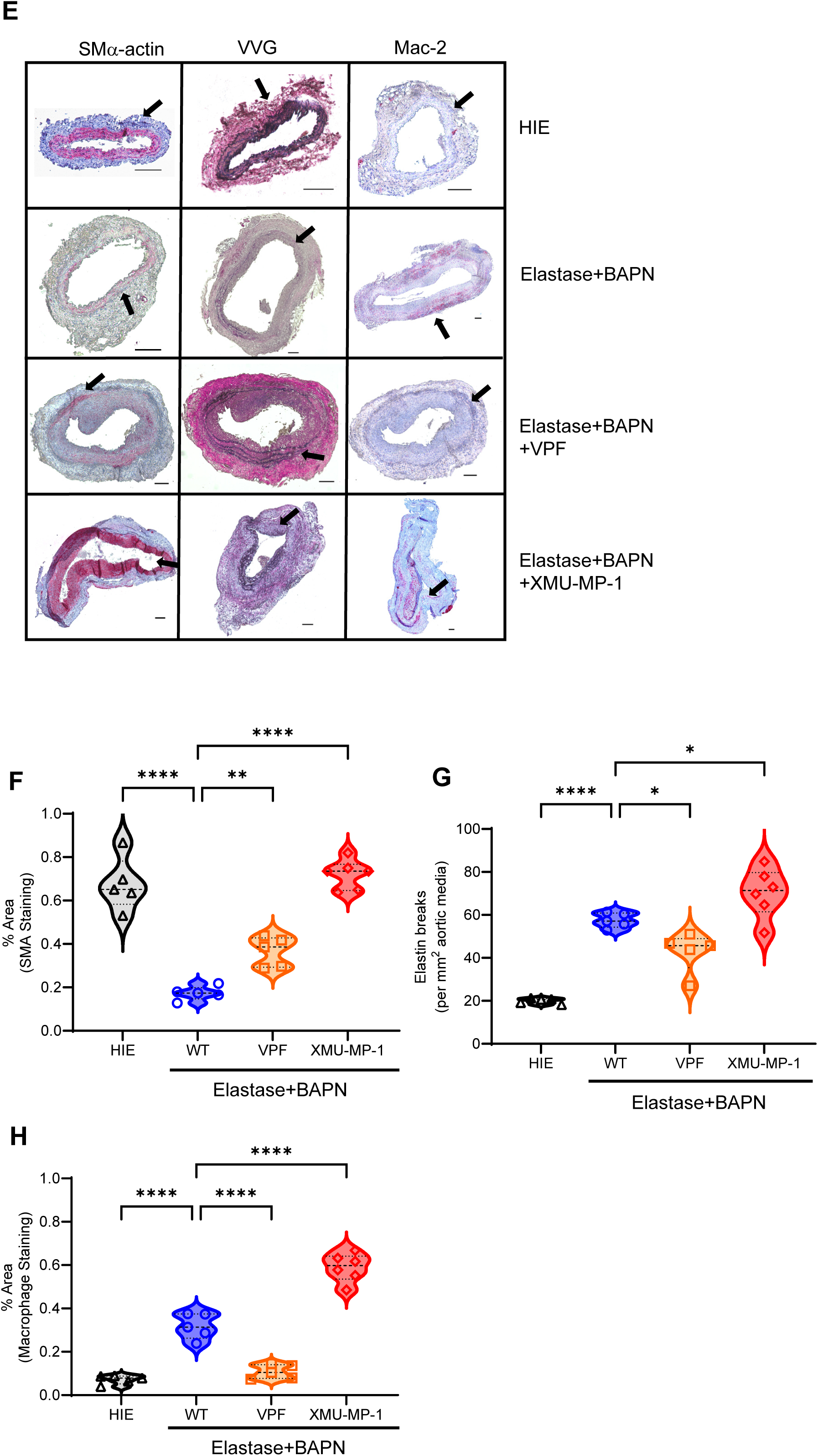
Verteporfin treatment mitigates aortic inflammation and vascular remodeling in a chronic AAA and rupture murine model. **A,** Schematic description of a 28-day chronic AAA and rupture topical elastase+BAPN with C57BL/6 mice. Male WT mice were induced with topical elastase two days after starting 0.2% BAPN water and injected with vehicle control, VPF, or XMU-MP-1. **B,** Verteporfin significantly reduced AAA diameter compared to elastase+BAPN treatment alone. Conversely, XMU-MP-1 significantly increased AAA size compared to elastase+BAPN-treated mice. ****p<0.0001, **p<0.01; n=12-23/group; 1-way ANOVA. **C,** Representative images of aortic phenotype in all groups. Scale bar=500μM. **D,** Kaplan Meier survival curve demonstrating a significant reduction in survival in XMU-MP-1 treated mice compared to elastase+BAPN and heat-inactivated controls. **E-H,** Comparative histology and quantification demonstrated preserved SM α-actin expression, decreased elastin fiber disruption, and reduced macrophage infiltration in VPF-treated mice compared to mice treated with elastase+BAPN. Conversely, XMU-MP-1 also showed increased α-SMA expression, but with increased elastin fiber breakage, and macrophage staining compared to mice treated with elastase+BAPN alone. Scale bar=100μM; ****p<0.0001, **p<0.01, *p<0.05; n=5/group; 1-way ANOVA.

Administration of VPF preserved SMα-actin expression (36.6±3.1% vs. 17.3±1.4%; p<0.01), decreased elastin fiber disruption (42.9±4.1 vs. 57.5±1.5; p<0.05), and reduced macrophage infiltration (10.8±1.4% vs. 31.8±2.6%; p<0.0001) compared to mice treated with elastase+BAPN on comparative histology and immunohistochemistry (Figure 4E-H). However, mice treated with XMU-MP-1 demonstrated increased SMα-actin expression (72.1±2.8% vs. 17.3±1.4%; p<0.0001), elastin fiber breaks (70.3±4.7 vs. 57.5±1.5; p<0.05), and macrophage infiltration (58.9±2.7% vs. 31.8±2.6%; p<0.0001) compared to mice treated with elastase+BAPN alone. These phenotypic and histologic findings demonstrated the ability of YAP/TAZ inhibition to mitigate AAA dilation and impending aortic rupture.

### Verteporfin treatment YAP-dependent remodeling and mechanotransduction pathways in ECs

To evaluate the EC-specific mechanistic role of YAP/TAZ modulation, human aortic endothelial cells (HAECs) were used for bulk RNA sequencing that revealed distinct gene profiles (Figure 5A). Elastase-treated HAECs exhibited an upregulation of differentially expressed genes involved in TNF/NF-kB activation (TNFAIP3, TRAF1, BIRC3), leukocyte recruitment and adhesion (VCAM1, ICAM1, SELE, CCL5, CXCL2, CXCL3, CXCL5) and matrix remodeling (MMP10) when compared to control HAECs (Figure 5B and Supplementary Table S2). Importantly, HAECs treated with VPF treatment showed significant downregulation of differentially expressed genes involved in ECM remodeling (LTBP1, VCAN, MMP16, NOX4), pathologic angiogenesis (KDR), and mechanotransduction (ITGA2, ITGA6), while HAECs treated with XMU-MP-1 exhibited upregulation of genes involved in cellular stress responses (RASD1, ZFP36), metabolic remodeling (CKB), and immune modulation/chronic inflammation (TNFSF9), compared to elastase-treated HAECs (Figure 5C-D and Supplementary Table S2). Together, these findings suggest that VPF and XMU-MP-1 act on ECs to downregulate vascular remodeling and mechanotransduction pathways, or promote pathways associated with endothelial-stress response, immune crosstalk, and vascular inflammation, respectively.

**Figure 5.**
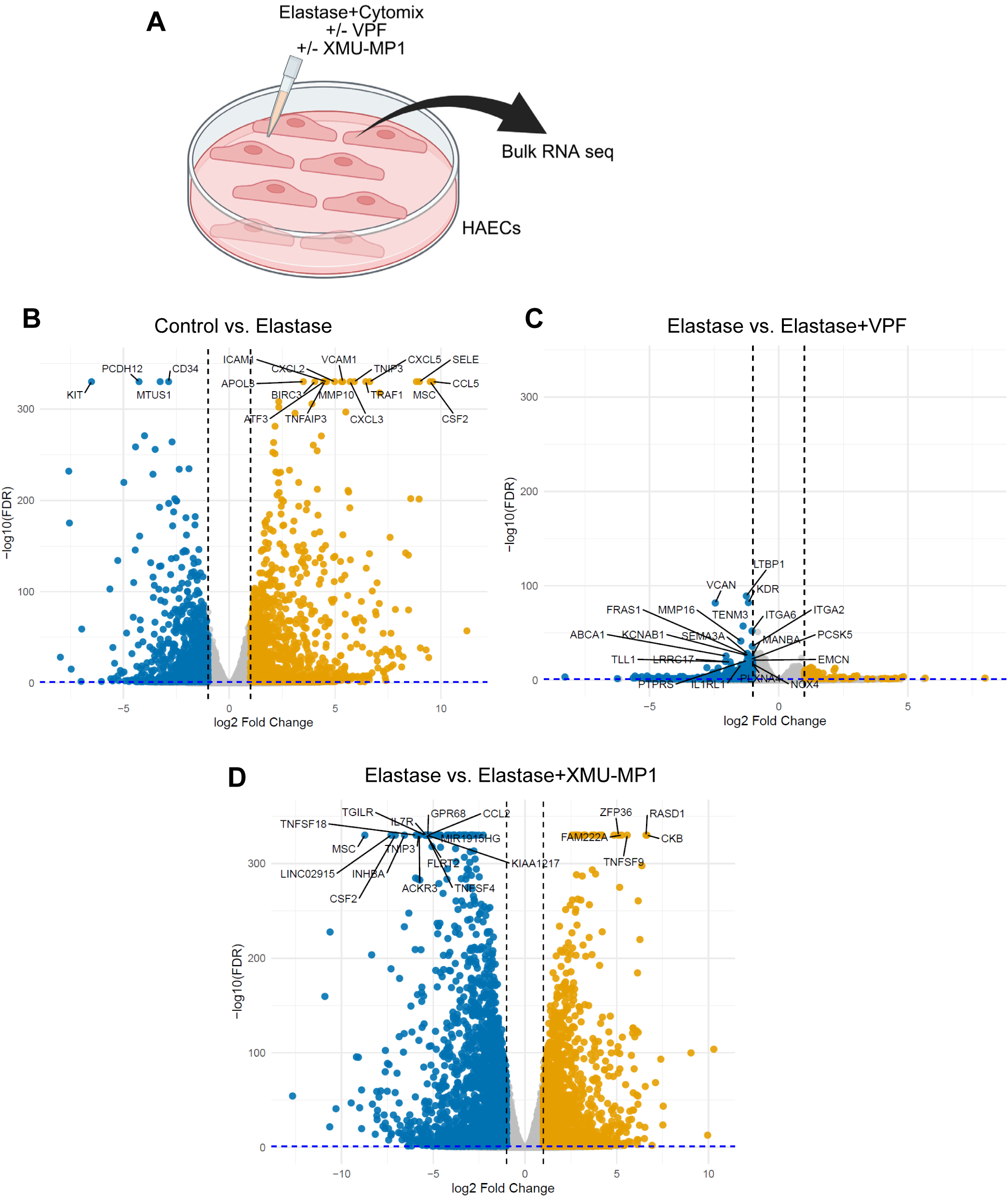
Verteporfin inhibits vascular remodeling in ECs. **A,** Schematic representing human aortic endothelial cells (HAECs) treated with elastase+cytomix (IL-17A+HMGB1+TNF-α; 10ng/ml), with or without VPF or XMU-MP-1 treatments. **B,** Volcano plot of differentially expressed genes in HAECs treated with elastase vs. control. Elastase-treated HAECs showed a significant upregulation of differentially expressed genes involved in TNF/NFkB activation, leukocyte recruitment and adhesion, and matrix remodeling when compared to control HAECs. **C,** Volcano plot of differentially expressed genes in HAECs treated with elastase+VPF vs. elastase. VPF treatment resulted in significant downregulation of differentially expressed genes involved in ECM remodeling, angiogenesis, and mechanotransduction compared to elastase-treated HAECs. **D,** Volcano plot of differentially expressed genes in HAECs treated with elastase + XMU-MP-1 vs. elastase. XMU-MP-1 treatment resulted in significant upregulation of differentially expressed genes involved in cellular stress response, metabolic remodeling, and immune modulation/chronic inflammation when compared to elastase-treated HAECs.

**Figure 6.**
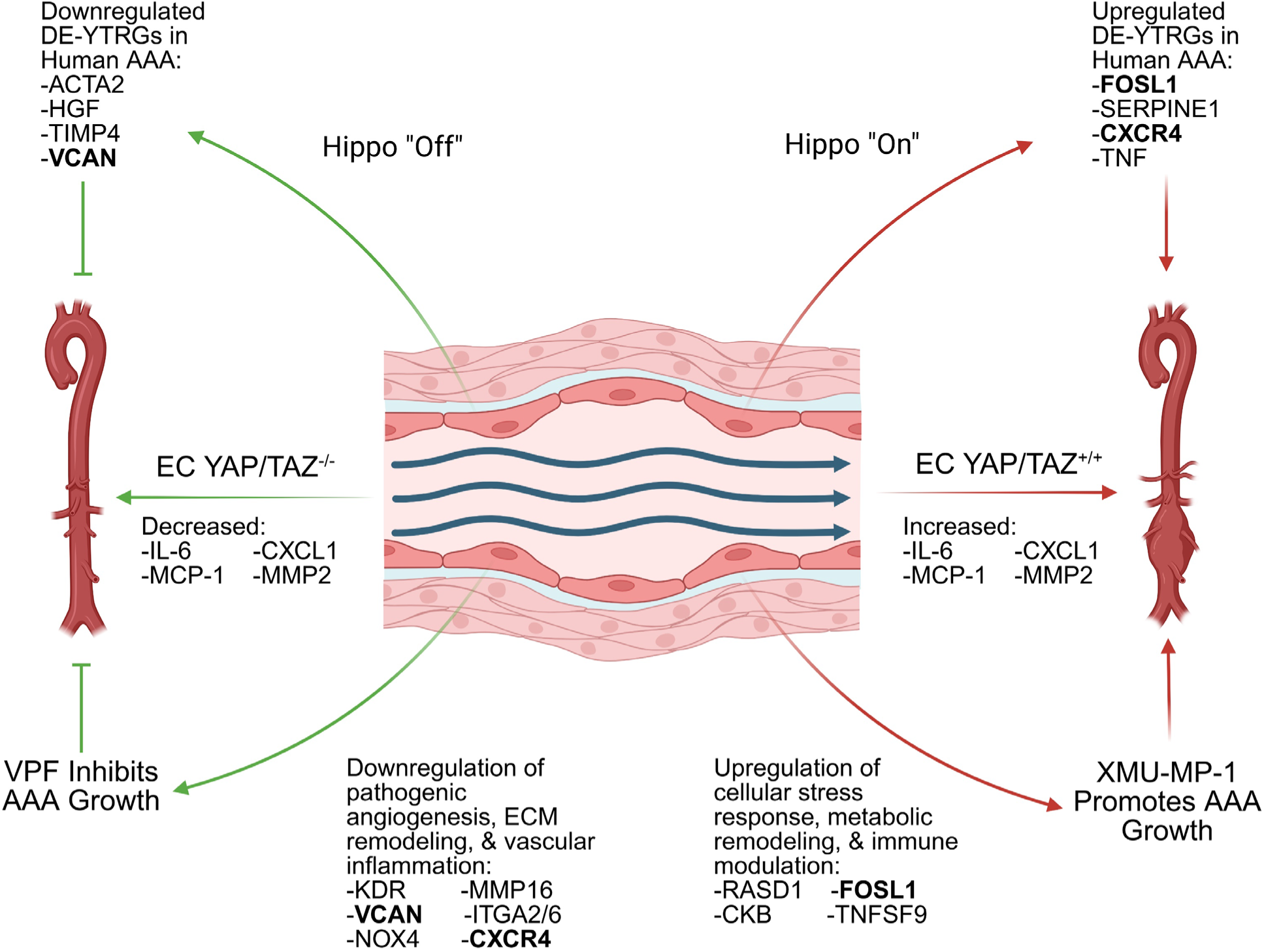
YAP/TAZ signaling on ECs inhibits AAA growth by downregulating ECM remodeling, pathologic angiogenesis, and mechanotransduction pathways. EC-specific YAP/TAZ deletion results in preservation of aortic morphology, decreased expression of pro-inflammatory cytokine expression, and smaller aneurysms. Treatment with VPF leads to downregulation of genes involved in ECM remodeling, pathologic angiogenesis, and inflammation. Conversely, activation of Hippo signaling via XMU-MP-1 promotes upregulation of genes involved in cellular stress response, metabolic remodeling, and immune modulation to promote AAA growth and rupture. DE-YTRGs, differentially expressed YAP/TAZ-related genes; ECM, extracellular matrix; VCAN, versican; FOSL1, Fos-related antigen 1; CXCR4; C-X-C motif chemokine receptor 4.

## DISCUSSION

This current study defines the importance of endothelial-cell specific YAP/TAZ activation in AAA formation. Our results show that YAP/TAZ mRNA and protein expression are increased in human AAA and specifically dysregulated at the endothelial cell level. Furthermore, we demonstrate that EC-specific YAP/TAZ expression increases sequentially during experimental AAA, and specific deletion of YAP/TAZ in ECs protects against AAA by mitigating aneurysm size, aortic remodeling, and pro-inflammatory cytokine and chemokine expression. Pharmacologic YAP/TAZ inhibition with VPF attenuates aneurysm formation, while activation via XMU-MP-1 exacerbates aortic growth, inflammation, leukocyte infiltration, and risk of rupture in two separate models of murine AAA. *In vitro* experiments confirm that pharmacologic modulation of YAP/TAZ occurs at the endothelial cell level, with bulk-RNA seq results suggesting a vasculoprotective effect of VPF through suppression of YAP-dependent remodeling and mechanotransduction, while cells treated with XMU-MP-1 exhibit an upregulation of pathways associated with EC dysfunction and pathological vascular remodeling.

AAAs are characterized by medial degeneration, elastin fragmentation, and chronic inflammation mediated by matrix degrading enzymes and the increased production of pro-inflammatory cytokines.^6–8^ The cumulative effect of this chronic inflammatory process leads to a progressive weakening and dilation of the aortic wall with an increasing risk of rupture and death.^4,5,43^ Recent reports suggest that the transcriptional co-activators YAP and TAZ play an important in aneurysm pathogenesis, vascular aging, and cardiovascular mechanotransduction signaling pathways.^15,44^ These central mediators of the Hippo pathway are highly sensitive to biomechanical stress, cytoskeletal tension, and extracellular matrix stiffening.^17^ Under physiologic laminar flow conditions at the endothelial cell level, YAP and TAZ are phosphorylated by a series of upstream Hippo kinases, including MST1/2 and LATS1/2, and deactivated in the cytoplasm for subsequent degradation. However, during oscillatory flow conditions, the Hippo pathway is turned off and YAP/TAZ are allowed to translocate to the nucleus to induce cell proliferation, angiogenesis, and atheroprone pathways.^45–47^

Recent studies examining the effects of YAP/TAZ on aging in *Fbn1* mutant mice noticed that inducing an active form of YAP in vascular smooth muscle cells protected against medial and adventitial thickening and elastin degradation.^14^ The role of Hippo signaling in other vascular cells show that deletion of YAP/TAZ in SMCs led to development of spontaneous aneurysms characterized by elastin disarray, apoptosis, and accumulation of pro-inflammatory immune cell populations.^11^ These inflammatory effects were mediated via cGAS/STING pathway during loss of YAP/TAZ in vascular smooth muscle cells. To date, however, there is very limited evidence for the role of EC-specific YAP/TAZ in AAA pathogenesis.

As ECs have been shown to play an important role in AAA pathogenesis and inflammatory milieu, we sought to elucidate the contributions of EC-specific YAP/TAZ to aortic inflammation and vascular remodeling in AAA formation. Previous studies have examined the role of YAP/TAZ activation in pathological angiogenesis and vascular inflammation, citing its role in leukocyte-endothelial adhesion via VCAM1 expression in the setting of TNFα secretion and stretch-induced proliferation via VE-Cadherin and Notch signaling during angiogenesis.^48–50^ Our scRNA seq analysis identified 242 differentially expressed YAP/TAZ-related genes that were upregulated in AAA tissue compared to controls. Of note, these genes included FOSL1, SERPINE1, and CXCR4, which have been shown to play a role in aneurysm pathogenesis. FOSL1 encodes a subunit of the activator protein 1 (AP-1) complex, which plays a critical role in cell proliferation and angiogenesis and has been found to mediate collagen synthesis and myofibroblast transformation in angiotensin-II murine models.^51^ Similarly, overexpression of SERPINE1 and CXCR4 have been shown to lead to vascular stiffening, enhanced endothelial cell apoptosis, and acceleration of aneurysm progression and rupture in humans and murine models of AAA and dissection.^52–56^ We also identified differentially expressed YAP/TAZ-related genes that were downregulated in human AAA, including ACTA2, HGF, and SYNPO2. ACTA2, which encodes α-SMA, and SNYPO2 are key structural genes involved in maintaining the stability of smooth muscle cells and the actin cytoskeleton.^57,58^ Decreased expression has been associated with medial degeneration, vessel wall instability, aneurysm formation, and vascular smooth muscle cell phenotypic switching in human AAA.^59,60^ Moreover, hepatocyte growth factor (HGF) plays a protective role in the development and rupture of aneurysms by promoting an anti-inflammatory cytokine profile.^61,62^ These differentially expressed YAP/TAZ related genes in AAA correspond to the phenotypic changes seen in our murine model of AAA, in which we show that EC-specific YAP/TAZ signaling mitigates aneurysm formation by reducing aortic remodeling through the preservation of SMA and elastin fiber integrity and blunting the inflammatory response through the reduction of macrophage infiltration and pro-inflammatory cytokine and chemokine expression.

Pharmacologic alteration of YAP/TAZ and the Hippo signaling pathway has garnered significant interest not only in aortic aneurysm pathophysiology, but also in aging and cancer. VPF is a benzoporphyrin photosensitizing agent that is FDA approved for photodynamic therapy in patients with age-related macular degeneration.^63^ Mechanistically, it acts as an inhibitor of YAP with its TEAD binding domain, blocking the transcriptional activation of downstream targets.^18,63,64^ Recently, Xie *et al*. discovered that blocking YAP with VPF decreased AAA incidence and collagen deposition in adventitial fibroblasts using a murine elastase model of AAA.^18^ Our murine and *in vitro* results corroborate these findings and suggest that VPF acts on ECs to suppress ECM remodeling, pathological angiogenesis, and mechanotransduction. *In vitro* bulk RNA seq results suggest that VPF acts on HAECs to ameliorate pathogenic angiogenesis, ECM remodeling, and vascular inflammation, as VPF-treated HAECs exhibited downregulation of differentially expressed genes including KDR, VCAN, NOX4, and ITGA2/6.^65–67^ KDR (kinase insert domain receptor) encodes vascular endothelial growth factor receptor 2, of which overexpression has been shown to lead to pathogenic angiogenesis and neovascularization in AAA.^67,68^ Data on VCAN is more limited to AAA formation, however, studies on the thoracic aorta have shown that accumulation of versican is associated with aneurysm formation.^66,69^ Moreover, our scRNA seq analysis revealed that VCAN is significantly differentially expressed in human AAA, suggesting that accumulation may promote pathologic remodeling. Finally, NOX4 and ITGA2/6 have been shown to promote oxidative stress and vascular inflammation via the production of reactive oxygen formation species, MMP expression, and cause vascular smooth muscle cell phenotypic switching seen in AAA formation.^70–73^

Additionally, there is limited research to date on the effects of the MST1/2 kinase inhibitor, XMU-MP-1, on AAA development and progression. Okuyama et al. found that Hippo-YAP signaling proteins were significantly elevated in angiotensin II-infused ApoE mice, and that treatment with XMU-MP-1 resulted in an equivocal increase in AAA diameter.^29^ Using two models of murine AAA, our data shows that XMU-MP-1 treatment exacerbates AAA formation and rupture, pro-inflammatory cytokine production, and aortic remodeling, suggesting that over-activation of the YAP/TAZ pathway can have a deleterious effect on AAA. XMU-MP-1 treatment of HAEC’s treated with elastase showed differentially expressed genes associated with cellular stress responses, metabolic remodeling, and chronic inflammation, including RASD1, CKB, and TNFSF9.^74–76^ Moreover, HAECs treated with XMU-MP-1 exhibited a significant upregulation of FOS1 – similar to analysis of human AAA scRNA seq. Together, our scRNA seq analysis, murine models of AAA, and bulk RNA seq results correlate with previous studies on the vasculoprotective effects of endothelial cell YAP/TAZ deletion and inhibition.^47,77,78^

A few limitations should be considered for this study. In addition to chemically induced AAA models used here, the role of YAP/TAZ as mechanosensitive transcription factors involved in vascular homeostasis and changes in oscillatory flow, should also be conducted using an angiotensin II/ApoE^-/-^ model to investigate the role of Hippo signaling in hypertension-induced AAAs. Moreover, our results point towards a vasculoprotective effect of VPF, while human aortic ECs treated with XMU-MP-1 exhibit an upregulation in pathways associated with endothelial dysregulation and impaired signaling. Further experiments with confirmatory qPCR and immunostaining of murine aortas should be conducted to delineate the mechanistic effects of our differentially expressed YAP/TAZ-related genes involved in sequential vascular remodeling.

In summary, we have shown that EC-specific YAP/TAZ signaling mediates aortic inflammation and vascular remodeling during AAA formation. Pharmacologic inhibition of YAP/TAZ offers a targetable therapeutic option to mitigate AAAs and prevent aortic rupture. Mechanistically, immunomodulation of mechanotranduction-related genes secondary to Hippo signaling downregulates endothelial-leukocyte crosstalk and mitigates aortic remodeling offering a restoration of vascular homeostasis.

## Acknowledgements

We thank Tabitha Randi for technical assistance in our laboratory.

## Supplemental Material

Tables S1-2

Figures S1-5

Major Resources Table

**Footnote**

## Nonstandard Abbreviations and Acronyms

AAA: abdominal aortic aneurysms
EC: endothelial cells
SMC: smooth muscle cells
VPF: verteporfin
MST: mammalian STE20-like protein kinase
LATS: large tumor suppressor
FOSL1: FOS like 1
SERPINE1: serpin family E member 1
CXCR4 C-X-C: motif chemokine receptor 4
ACTA2: actin alpha 2, smooth muscle
HGF: hepatocyte growth factor
SYNOP2: synaptopodin 2
BAPN: β-Aminopropionitrile
IL: interleukin
MCP-1: monocyte chemoattractant protein-1
CXCL1: chemokine (C-X-C motif) ligand 1
MMP: matrix metalloproteinase
scRNA-seq: single-cell RNA sequencing
YTRGs: YAP/TAZ-related genes
VVG: Verhoeff-van Gieson

