## Supplementary material for "YAP/TAZ Signaling in Endothelial Cells Mediates the Pathogenesis of Abdominal Aortic Aneurysm Formation": Supplemetal Figures S1-S5

### Supplementary Figure S1

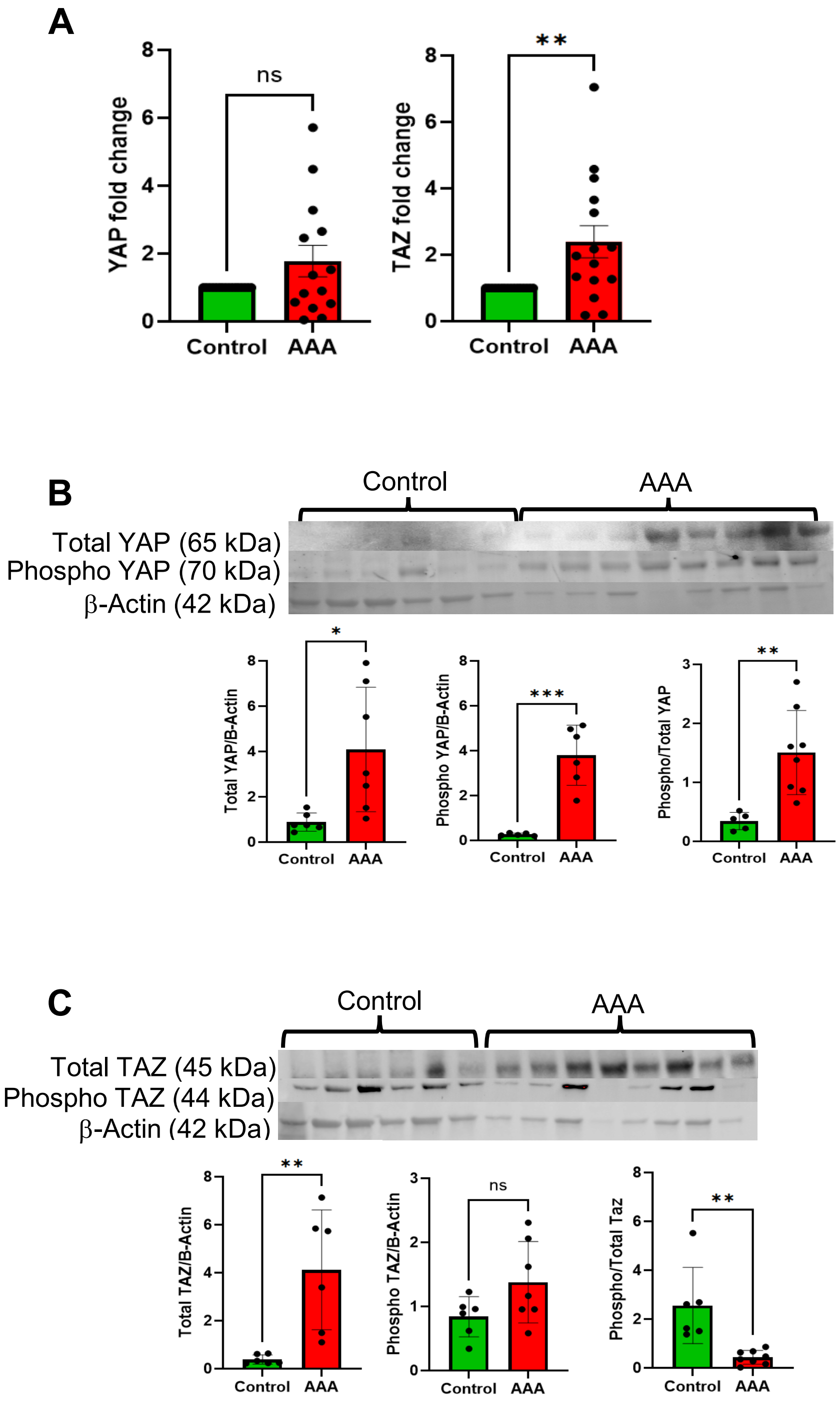


**Supplementary Figure S1**. **YAP/TAZ expression is altered in human AAAs**. **A**, Human AAA tissue showed a significant increase in TAZ mRNA compared to controls. n=14-16/group; **p=0.01; unpaired t-test. There was no significant change in mRNA expression for YAP. n=14-16/group; p=0.08; ns, not significant; unpaired t-test. **B-C**, Total YAP and TAZ protein expressions were significantly increased in AAA compared to controls. n=6-7/group; *p<0.05; **p<0.01; ***p<0.0001; unpaired t-test.

### Supplementary Figure S2

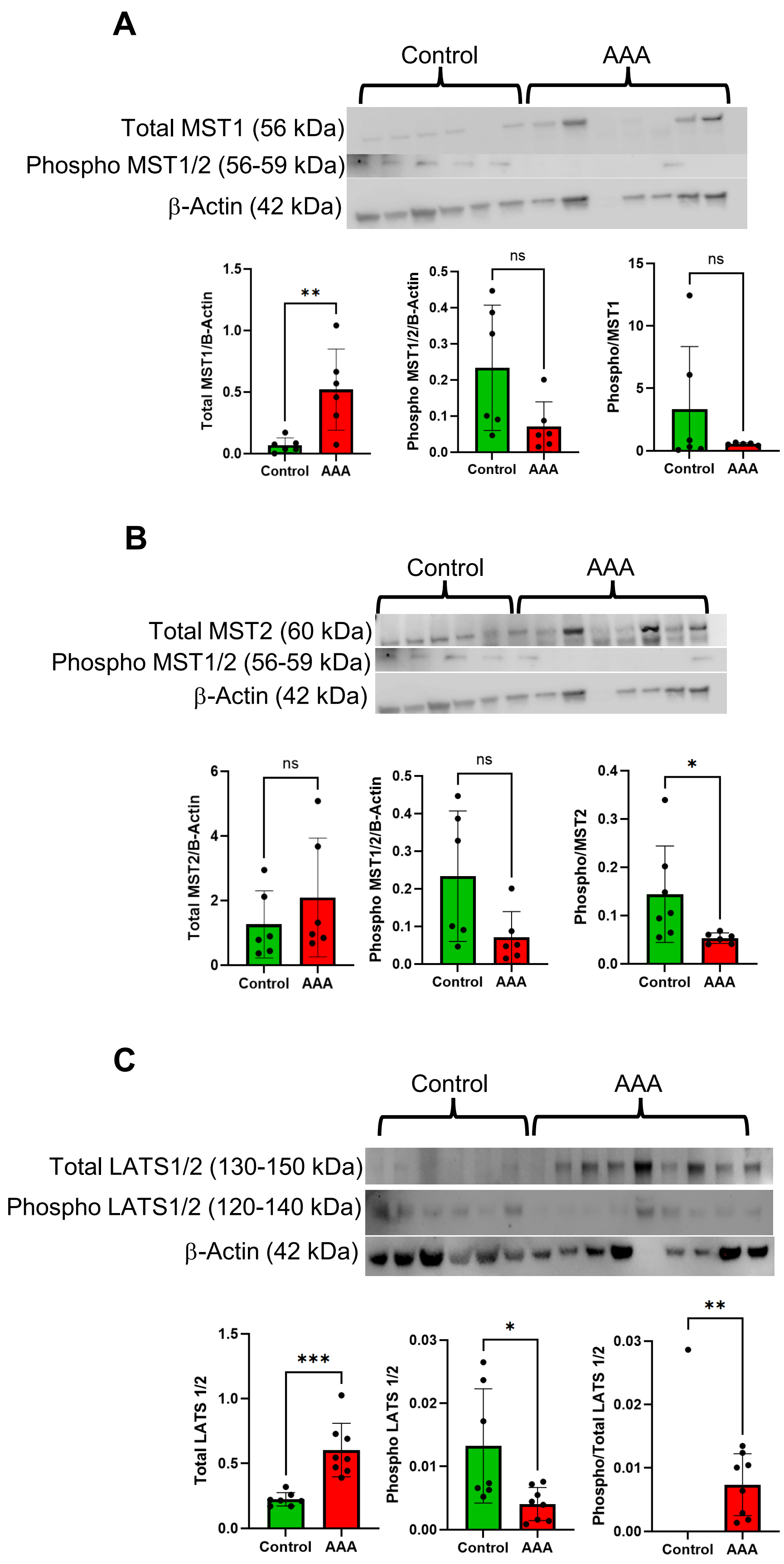


**Supplementary Figure S2**. **The Hippo pathway is downregulated during AAA promoting YAP/TAZ nuclear localization**. **A**, Human AAA tissue showed a significant increase in the upstream Hippo pathway kinase total MST1; n=6/group; **p<0.01; unpaired t-test. **B**, The phosphor/total MST2 expression ratio was significantly attenuated in AAAs compared to age-matched controls. n=6/group; ns; not significant; *p<0.05;unpaired t-test. **C**, LATS1/2 protein expression is also significantly increased in AAA tissue compared to controls. n=6/group; *p<0.05; **p<0.01; ***p<0.0001; unpaired t-test.

### Supplementary Figure S3

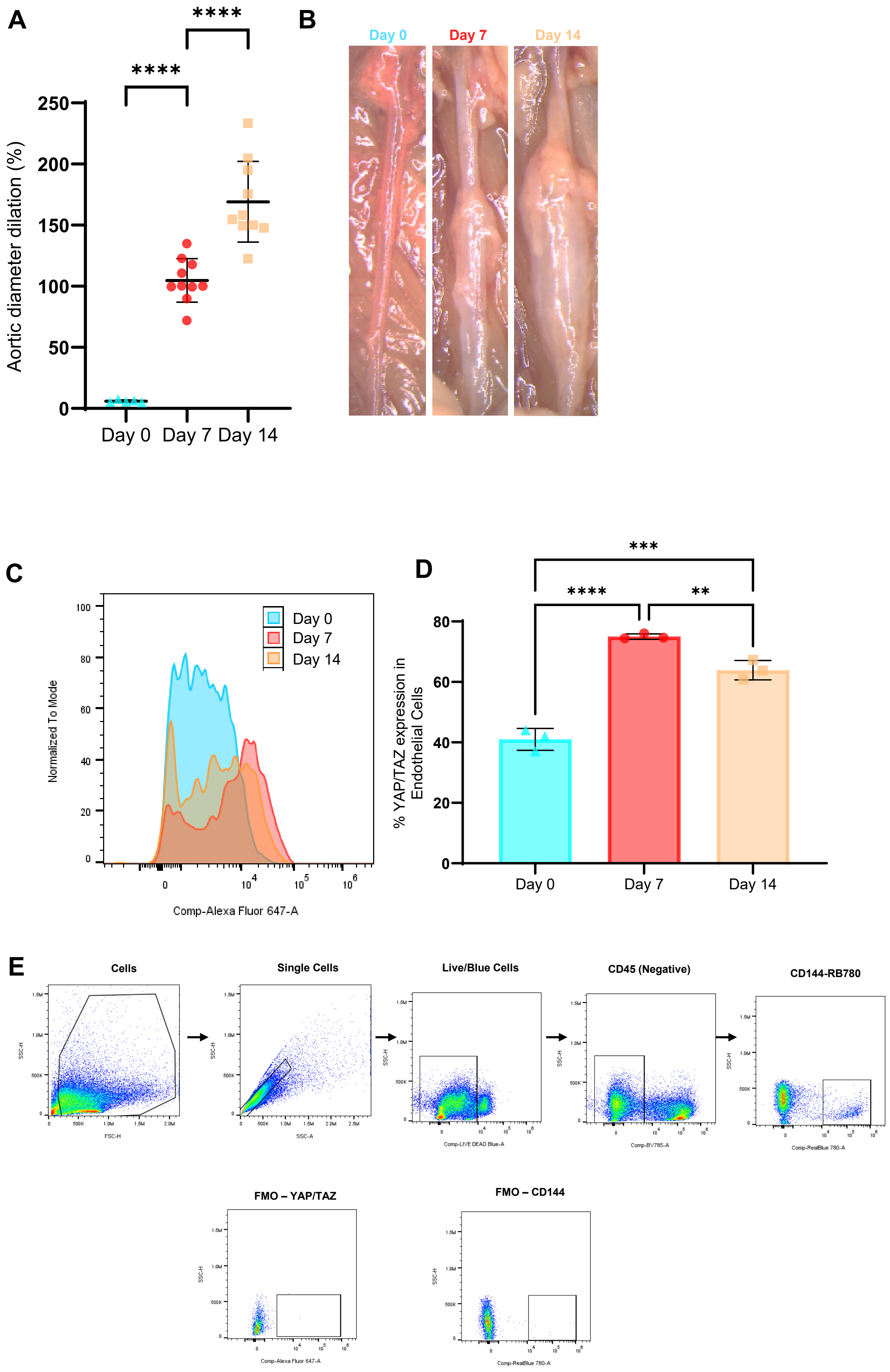


**Supplementary Figure S3. YAP/TAZ expression in ECs increases in murine AAAs. A,** Murine aortic diameter increases over time in the topical elastase AAA model, with maximal aortic diameter occurring on day 14. **B,** Representative images of aortic phenotype in all groups. Scale bar=500μM. **C-D,** Flow cytometry analysis revealed that YAP/TAZ expression in ECs increases over time, peaking at day 7 where aortic endothelial cells show increased YAP/TAZ expression compared to day 0. n=5-10/group; **p<0.01; ***p<0.0001; ****p<0.0001; 1-way ANOVA. **E,** Gating strategy to analyze EC-specific YAP/TAZ expression during murine AAA formation.

#### Supplementary Figure S4

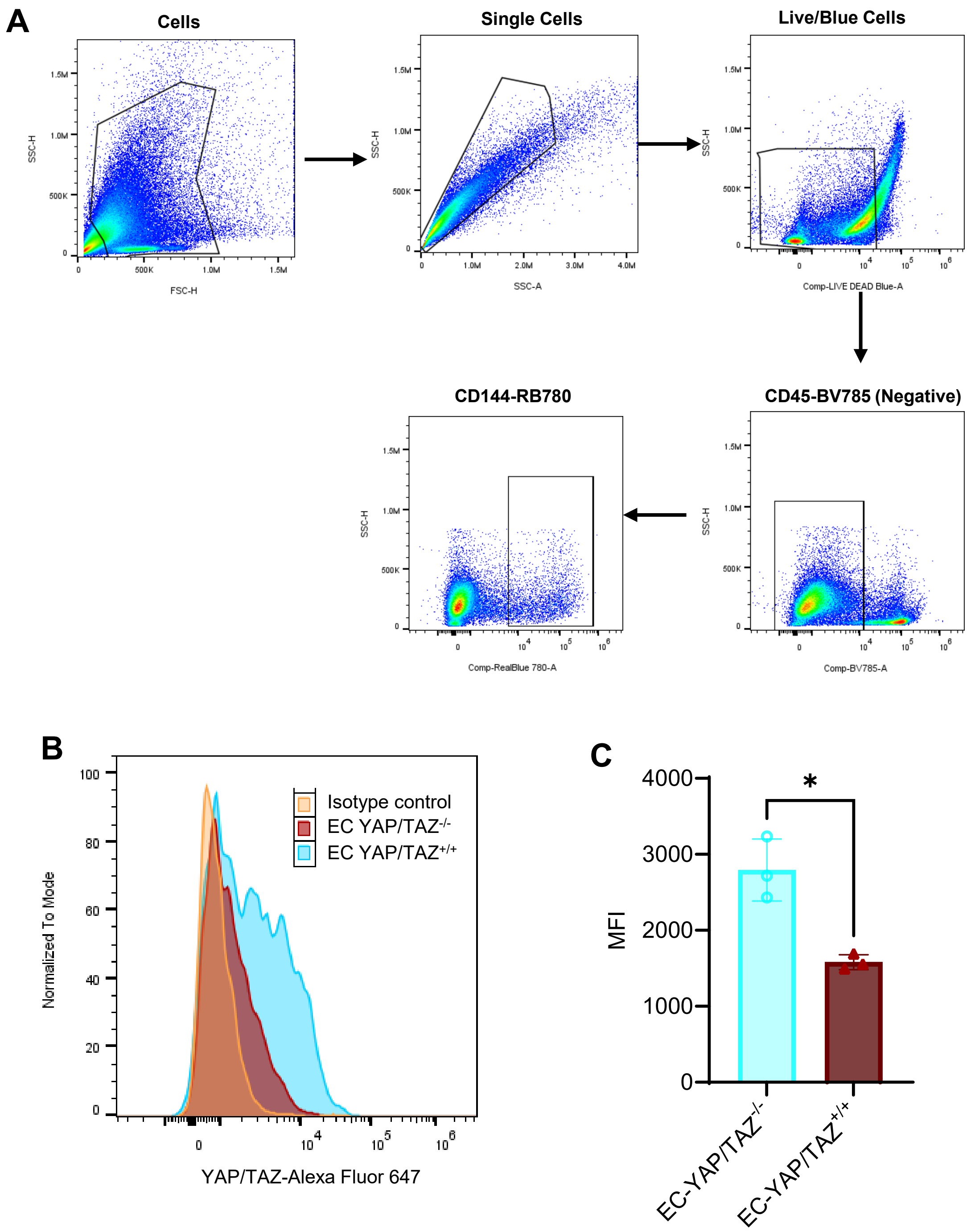


**Supplementary Figure S4. Confirmation of YAP/TAZ deletion in ECs by flow cytometry. A,** Gating strategy to confirm EC YAP/TAZ deletion. **B-C,** A significant decrease in YAP/TAZ expression was observed in EC YAP/TAZ$^{-/-}$ mice compared to littermate controls. n=3/group; *p<0.03.

#### Supplementary Figure S5

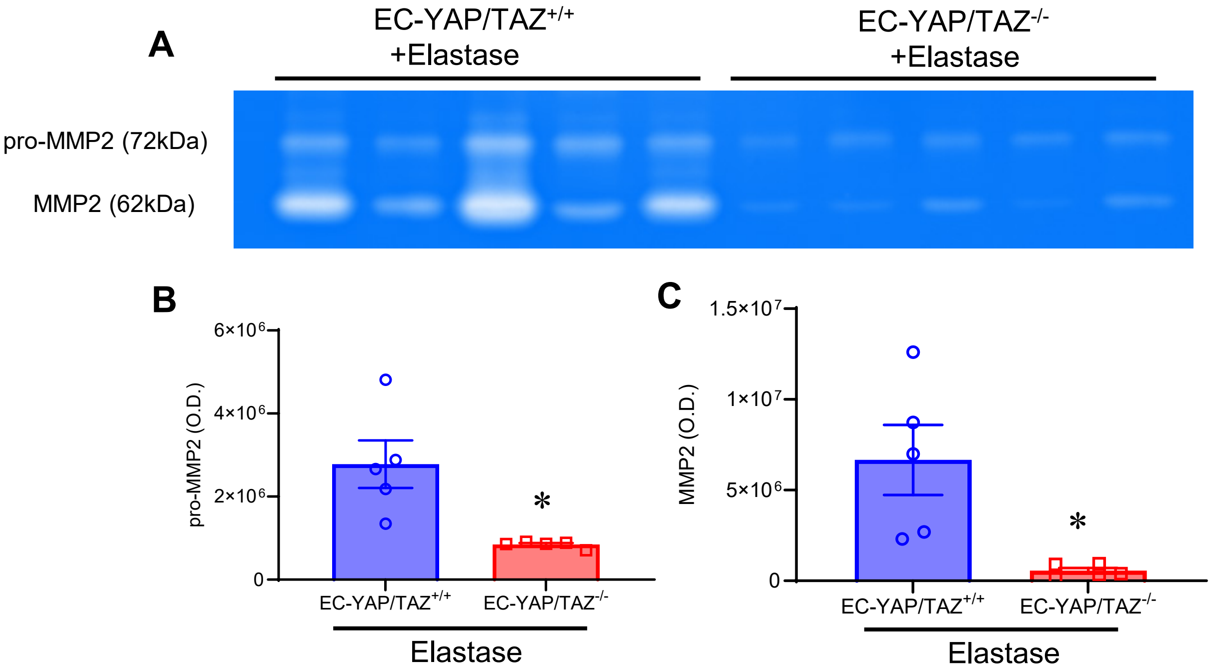


**Supplementary Figure S5. YAP/TAZ deletion in ECs reduces MMP2 expression. A,** MMP2 activity was measured in aortic tissue by zymography on day 14 using the topical elastase model of AAA. **B-C,** Quantification of zymograms demonstrates that elastase-treated EC-YAP/TAZ deletion significantly reduces MMP2 activity compared to elastase-treated littermate controls. n=5/group; *$p<0.008$; Mann-Whitney test..
